# Cancer subtype identification using somatic mutation data

**DOI:** 10.1101/228031

**Authors:** Marieke L. Kuijjer, Joseph N. Paulson, Peter Salzman, Wei Ding, John Quackenbush

## Abstract

With the onset of next generation sequencing technologies, we have made great progress in identifying recurrent mutational drivers of cancer. As cancer tissues are now frequently screened for specific sets of mutations, a large amount of samples has become available for analysis. Classification of patients with similar mutation profiles may help identifying subgroups of patients who might benefit from specific types of treatment. However, classification based on somatic mutations is challenging due to the sparseness and heterogeneity of the data.

**METHODS:** Here, we describe a new method to de-sparsify somatic mutation data using biological pathways. We applied this method to 23 cancer types from The Cancer Genome Atlas, including samples from 5, 805 primary tumors.

**RESULTS:** We show that, for most cancer types, de-sparsified mutation data associates with phenotypic data. We identify poor prognostic subtypes in three cancer types, which are associated with mutations in signal transduction pathways for which targeted treatment options are available. We identify subtype-drug associations for 14 additional subtypes. Finally, we perform a pan-cancer subtyping analysis and identify nine pan-cancer subtypes, which associate with mutations in four overarching sets of biological pathways.

**CONCLUSIONS:** This study is an important step towards understanding mutational patterns in cancer.

## BACKGROUND

Cancer is a heterogeneous disease that can develop in different tissues and cell types. Even within one cancer type, the disease may manifest itself in multiple subtypes, which are usually distinguished based on different histology, molecular profiles, or specific mutations, and which may lead to different clinical outcomes. Identifying new cancer subtypes can help classification of patients into groups with similar clinical phenotypes, prognosis, or response to treatment. As an example, breast cancer is typically classified into four primary molecular subtypes based on expression of *HER2*, hormone receptors, and tumor grade, and these different subtypes have different prognosis and respond differently to hormone therapy [1]. While these subtypes are used to manage patient treatment, even here we know that individual subtypes themselves represent a diversity of smaller groups.

Since the onset of large-scale genomic experiments, cancer subtypes have been identified in multiple cancers, using mRNA [2, 3] and microRNA expression levels [2], methylation data [2, 4], copy number alterations, and combinations of different ‘omics data types [5], but few studies have subtyped patients based on somatic mutations. Somatic mutations play a large role in cancer development and disease progression, and mutational profiling is used far more commonly than other ‘omics analyses in clinical practice because most clinical guidelines are based on single gene mutations. Consequently, classification based on patterns of mutation could be particularly informative for identification of subgroups of patients who might respond to specific targeted treatment regimens and of those who are unlikely to respond.

However, subtype classification using somatic mutations in cancer is challenging, mainly because the data are very sparse: many tumors only have a handful of mutations in coding regions yet the total number of mutations within a population is typically substantial. Often, frequent cancer drivers—such as *TP53*—are mutated, as well as so-called “passenger” events that are considered mutational noise yet which may still influence tumor properties. And even within the same cancer type, tumors often exhibit very different mutational patterns, including drivers and passengers—as well as mutations that may fall somewhere in between.

To classify sparse somatic mutation data into subtypes, published methods generally first de-sparsify the data. Some methods use a gene-gene network as “prior” knowledge to de-sparsify the data [6–9]. Hofree *et al.* [7], for example, use network propagation to “fill in” the mutational status of neighboring genes (in protein-protein interaction networks) of mutated drivers, while Le Morvan *et al.* [9] use networks from Pathway Commons to normalize a patient’s mutational profile by adding “missing” or by removing “non-essential” mutations.

Data de-sparsification using gene-gene networks has been helpful in identifying subnetworks involved in cancer [8], as well as in identifying genes associated with patient survival [9]. However, gene-gene networks depend on a set of known “prior” interactions, but these priors may or may not be “correct” in the sense that they may not be relevant to the tissue or tumor under study. This reliance on “canonical” networks might over-emphasize genes that are connected to mutational drivers through such interactions, as well as over-emphasize highly connected genes, even though some studies do correct for this [8].

In addition, genes belonging to the same biological pathway do not necessarily have to be closely linked in a gene-gene (protein-protein interaction) network—they do not always interact physically, and their functional interactions may be indirect. However, having multiple mutations in the same biological pathway is likely disruptive to the pathway’s function, and likely more so than only having one gene mutated in that pathway. Thus, to classify somatic mutation data into meaningful subtypes, we believe it is important to take all genes in a pathway into account when de-sparsifying the data.

Finally, because somatic mutation data are very heterogeneous, pan-cancer studies may help understanding the biological processes that play a role in cancer. The inclusion of multiple cancer types in an analysis both increases the sample size and allows for the discovery of mutation subtypes across cancer types. Large consortia, including The Cancer Genome Atlas (TCGA) and International Cancer Genome Consortium (ICGC), have performed comprehensive pan-cancer analyses of somatic mutations [10, 11]. Hoadley *et al.* [12] used pathway scores to integrate somatic mutations with other ‘omics data types to perform a multi-platform classification of twelve cancer types, while Leiserson *et al.* [8] identified pan-cancer subnetworks across the same twelve types of cancer. While these studies have improved our understanding of the genes and pathways that are recurrently mutated in cancer, data are now available for many more samples and cancer types, increasing the power to detect new mutational patterns and cancer subtypes.

In this study, we describe SAMBAR, or Subtyping Agglomerated Mutations By Annotation Relations, a method to de-sparsify somatic mutation data by summarizing these data into pathway mutation scores. We applied SAMBAR to data from 5, 805 primary tumors from TCGA, including 23 different cancer types. We used the de-sparsified data to associate mutational patterns with phenotypic data, to identify prognostic subtypes, and to identify potential drug targets associated with subtypes in each of the cancer types. In addition, we performed a pan-cancer analysis to identify mutation subtypes across multiple cancers, and describe the mutational patterns associated with these subtypes.

## METHODS

### Curation of clinical data

We used RTCGAToolbox [13] to download clinical data for 23 cancer types from TCGA. We curated these data by combining data from all available Firehose versions (accessed July 17, 2015) for each cancer type consecutively as to retain all clinical information, with the most up-to-date information for data present across different Firehose versions (Supplemental Materials and Methods, Supplemental Table 1).

**Table 1:** Drug targets enriched for mutations in cancer-specific subtypes. *k*: the level at which the subtyping dendrogram was cut, Patients: the number of patients in a subtype, versus the number of patients of that cancer type in our dataset, Enr.: enrichment, or the observed over expected ratio, Drugs: drugs of which targets were enriched for mutations in the subtype.

| Cancer | *k* | Patients | Enr. | Drugs |
| --- | --- | --- | --- | --- |
| ACC | 2 | 12/90 | 41.4 | idasanutlin, nutlin-3 |
| BLCA | 2 | 25/130 | 6.6 | amuvatinib, BMS-817378, cabozantinib, golvatinib, OSI-930, PD-153035, PLX-4720, ZM-39923 |
| BRCA | 2 | 16/956 | 7.6 | SGC-CBP30 |
| GBM | 2 | 62/280 | 5.4 | amuvatinib, BMS-817378, cabozantinib, golvatinib, lestaurtinib, OSI-930, PLX-4720 |
| HNSC | 2 | 69/305 | 4.7 | amuvatinib, AS-703026, atiprimod, AZ-628, AZD1480, baricitinib, CEP-32496, curcumol, DCC-2618, LY2784544, LY3009120, MEK162, PD-198306, peficitinib, refametinib, Ro-5126766, ruxolitinib-(S), TG-101209, trametinib, vemurafenib, XL019, ZM-39923 |
| KIRC | 2 | 19/419 | 9.2 | AG-490 |
| KIRP | 2 | 16/164 | 4.8 | bortezomib |
| LAML | 2 | 23/181 | 13.2 | lonafarnib |
| LGG | 2 | 36/287 | 12.6 | atiprimod, LY2784544, TG-101209 |
| LUSC | 2 | 28/177 | 6.4 | ZM-39923 |
| OV | 3 | 13/445 | 3.3 | bortezomib |
| PRAD | 2 | 13/239 | 20.7 | SGC-CBP30 |
| READ | 2 | 49/116 | 7.4 | PD-153035 |
| THCA | 2 | 284/357 | 5.6 | lonafarnib |
| UCS | 2 | 24/57 | 4.3 | wortmannin |

### Processing of mutation data

We downloaded .maf files (*n* = 47) containing mutation information for 6, 406 samples from 23 cancer types from the TCGA website (accessed March 17-18, 2014). We removed silent mutations and only retained genes with hg18/19 annotations (19, 065 genes). We divided the number of non-silent mutations *N_ij_* in a sample *i* and gene *j* by the gene’s length *L_j_*, defined as the number of non-overlapping exonic base pairs, calculated on either hg18 or hg19, depending on the annotation the sample was mapped to. We next removed samples that were not obtained from primary tumors. Finally, we merged replicate tumor samples that were derived from the same patient by taking the maximum mutation score for each gene, so that we retained all mutations that were observed in the tumor. The resulting dataset contained gene mutation scores for 5, 992 patients.

We further subsetted these data to 2, 219 cancer-associated genes from COSMIC [14] and Supplemental Table 3 from Östlund *et al.* [15]. For each patient, we calculated the overall cancer-associated mutation rate by summing up mutation scores in these genes *j^’^*. We removed samples with a rate of 0 (*n* = 108). For the remaining samples, we divided the mutation scores by these mutation rates, resulting in mutation rate-adjusted scores (*G*, Equation 1). We also removed samples which did not have mutations in the de-sparsified data (see below).

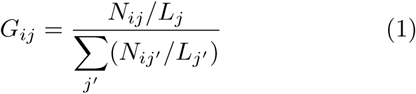

### De-sparsifying mutation data using biological pathways

We downloaded the file “c2.cp.v5.0.edges.gmt” from MSigDb, which included 1, 135 canonical pathway gene signatures (*q*). We converted this file into a binary matrix *M*, with information of whether a gene *j* belongs to a pathway *q*. We calculated pathway mutation scores (*P*) by correcting the sum of mutation scores of all genes in a pathway for the number of pathways *q^′^* a gene belongs to, and for the number of cancer-associated genes present in that pathway (Equation 2). We removed samples without mutations in any pathway (*n* = 79) and pathways without mutations in cancer-associated genes (*n* = 69), leaving us with 5, 805 patients and 1, 066 pathways for our subtyping analysis.

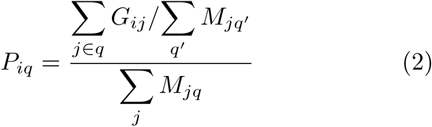

### Selecting the most variable distance metric

We used 12/16 distance metrics from the vegdist function in R package vegan [16] on both gene and pathway mutation scores to determine the metric that best separated patients. We excluded the “cao,” “chao,” “mountford,” and “morisita” metrics, because these are intended to be used on count data (integers) only. For each dataset and cancer type, we removed columns and rows that did not have any mutations prior to calculating distances between the samples.

### Correlating clinical variables to principal components

We calculated distance matrices of individuals within each cancer type using the binomial dissimilarity index on pathway mutation scores. We regressed each of the top five principal coordinates from the PCA on these distance matrices with each clinical variable that had ≥ 2 different entries. We generated word clouds using https://www.wordclouds.com/. As input we included parameters associated (nominal *p <* 0.01) with a top five principal component. We normalized this list to the number of times a variable was available across the 23 cancer types. We adjusted for multiple testing using Benjamini and Hochberg’s method [17] across each cancer and principal component to report significant associations in Supplemental Table 2.

### Association of pathway mutation scores with survival data

For each cancer type, we clustered the pathway mutation scores using hierarchical clustering with binomial distance. We cut the dendrogram at *k* = 2−4, removing clusters of size *<* 10. We used the log-rank test (*p <* 0.01) to identify significant differences between overall survival profiles, choosing the lowest *k* if multiple *k*’s resulted in significant prognostic subtypes for the same cancer type.

We ran 10, 000 sample label permutations for each of the prognostic subtypes, fixing *k* and subtype sample sizes to correspond to those of the subtypes we had identified. We then estimated the permutation p-value, defined as the fraction of permutations with p-values smaller than that from the log-rank test on the actual prognostic subtypes. We assigned a p-value of ≤ 1/10, 000 to subtypes for which all permutation p-values were higher than that of the original log-rank test. We corrected these p-values for multiple testing using the Benjamini-Hochberg method [17].

To determine what biological pathways drive the poor prognostic subtypes, we selected pathways that were mutated in *>* 90% of samples in the poor prognostic subtype and in *<* 50% in the subtypes with better prognosis.

### Integration of subtypes with drug targeting information

We scanned each tumor type for 2−4 subtypes, as described above (Methods). We selected signatures mutated in *>* 95% of samples belonging to a particular subtype and in *<* 10% of samples belonging to the remaining subtypes. We then selected all genes belonging to these signatures, and, for each gene, calculated the fraction of samples with mutations in the subtype of interest, and the same fraction in the remaining samples. We selected those genes that were more frequently mutated in the subtype of interest.

We downloaded drugs and their target genes from CMap [18] (accessed October 17, 2017). For each drug, we calculated how many of its known targets overlapped with this list of more frequently mutated, pathway-associated genes. We removed interactions with *<* 2 subtype-specific drug targets. Next, we calculated an “observed” score based on the number of subtype-specific targets divided by the total number of known targets for that drug. We also calculated an “expected” score based on the number of cancer-associated genes present in all 1, 066 pathways that were also present as drug targets in CMap (a total of 290 genes). We defined significant subtype-drug associations as those interactions that had enrichment scores (the observed/expected ratio) *>* 3.

Finally, we filtered this set of subtype-drug interactions by removing redundant subtypes—subtypes identified using a larger *k* that included the exact same set of patients as identified using a lower *k*.

### Identification of pan-cancer subtypes

We used binomial distance to cluster pathway mutation scores of all 5, 805 samples. We cut the clustering dendrogram at *k* = 2-1,000, filtering out clusters of size *<* 50. We observed several breakpoints which the largest subtype was split into new subtypes. We selected the highest *k* at such a breakpoint, for which *>* 90% of all samples were assigned to subtypes (*k* = 169), resulting in 9 pan-cancer subtypes). We used Fisher’s exact test to identify whether these subtypes were enriched (estimate *>* 4 and Bonferroni-adjusted p-value *<* 0.05) for particular cancer types.

We defined significantly mutated signatures as those signatures that were mutated in at least 95% of all samples in a pan-cancer subtype. We visualized average mutation scores of these pathways in a heatmap and identified four sets of pathways by row clustering (binomial distance) these data).

To make word clouds for these pathways, we identified the frequency of all 1, 356 unique words (separated by underscores in MSigDb) occurring in the 1, 066 pathways. We removed words that occurred *<* 3 times (1, 051 words). We then selected words belonging to one of the four sets of pathways, and calculated their observed frequency by dividing the number of times the word occurred in the set of pathways by the total number of words in that set. Next, we calculated the expected frequency by dividing the number of times the specific word occurred in all pathways by the total number of words in all pathways. We multiplied the observed/expected ratio by 10, rounded the number to an integer, and used that number of words as input for https://www.wordclouds.com/.

### Validation of pathway activation using protein abundance data

We validated our pan-cancer subtypes using orthogonal evidence on pathway activation. We downloaded Reverse Phase Protein Array (RPPA) data from TCGA using R package RTCGA.RPPA [19] (accessed February 21, 2018) and filtered these data for primary tumors (5, 790 patients, including 3814 patients which we had subtyped based on mutation data) and proteins that were available across all samples (121 proteins). We curated a protein activation signature for our “Set 1” subtype by selecting all available protein products (*n* = 15) of genes in the “Reactome PI3K/AKT activation” pathway, and a protein activation signature for our “Set 2” subtype based on protein products (*n* = 6) of genes in the “Reactome p53-Dependent G1 DNA Damage Response” pathway. We next calculated protein activation scores by summing up protein abundance levels for each of these signatures. We then performed a t-test between protein activation scores of “Set 1” in patients belonging to “Set 1” subtypes (S1–2) and other patients (S3–9), and between scores of “Set 2” pathways in patients belonging to “Set 2” subtypes (S4, S6, S8) and other patients (S1–3, S5, S7, S9) to identify significant differences (p-value *<* 0.05) in protein activation between the pan-cancer subtypes.

### Validation of pan-cancer subtype drug response

We downloaded variants detected using Whole Exome Sequencing from the Cancer Genome Project (CGP) (file “WES variants.xlsx” from http://www.cancerrxgene. org/downloads, accessed February 21, 2018). This file included non-silent mutations for 1, 001 cell lines and 19, 100 genes. We de-sparsified these data using SAMBAR, as described above. We next downloaded IC50 scores for drugs targeting “Set 1” pathways by selecting the target pathway “PI3K/MTOR signaling” (21 unique drugs) and for drugs targeting “Set 2” pathways by selecting the target pathway “DNA replication” (11 drugs) using CGP’s “Data download” tool (https://www.cancerrxgene.org/translation/drug/download#ic50, accessed March 26, 2018).

We divided the cell lines into two groups: those that had mutations in all 94 “Set 1” pathways (*n* = 156), and those that did not (*n* = 845), and performed a t-test to identify significant differences in response to drugs targeting PI3K/MTOR signaling (Benjamini-Hochberg adjusted p-values *<* 0.05). We note that for some of the drugs acting on PI3K/MTOR, replicate measurements were available. For those drugs, we pooled data from the replicates. We repeated this analysis on cell lines that had mutations in all 38 “Set 2” pathways (*n* = 681) and those that did not (*n* = 320) and drugs targeting DNA replication. We note that we identified a relatively high number of cell lines with mutations in all “Set 2” pathways compared to the number we identified in primary tumors. We believe this number is high because cell lines have more mutations (median number of mutations in cell lines is 158 compared to 91 in primary tumors). In addition, genes involved in DNA replication pathways are often mutated in cell lines to help immortalization.

## RESULTS

### De-sparsification of cancer mutation data

We aimed to identify subgroups of cancer patients that might benefit from specific targeted therapies. We hypothesized that we could identify cancer subtypes both within specific cancers and across all cancer types, using information on gene mutation status. We curated clinical data for 23 cancer types from TCGA (see Methods and Supplemental Materials and Methods), and preprocessed mutation data from 5, 805 primary tumors comprising 23 cancer types from TCGA (see Methods and Supplemental Figure S1).

We calculated gene mutation scores by normalizing the number of non-silent mutations in a gene to the gene’s length. Even though silent mutations can potentially be cancer drivers [20–22], we assumed that most are passenger mutations caused by the background mutation rate, and thus removed such mutations to control for this. To further control for mutational noise, we subsetted these data to 2, 219 genes with either known roles in cancer, or with functional connections to such genes (cancer-associated genes). We found that the filtered data were very sparse and difficult to assign to subtypes (see Supplemental Figure S2).

We hypothesized that summarizing the gene mutation scores into biological pathway scores would help to deparsify the data, as well as help identifying subgroups of patients who might respond to specific drugs targeting those pathways. We therefore used SAMBAR to de-sparsify the data by calculating mutation scores for 1, 135 canonical pathways from MSigDb [23]. In short, for each pathway, we summed up mutation scores of all genes belonging to that pathway and corrected for the pathway’s gene set size and the number of times a gene was represented in the full set of pathways (see Methods). We then corrected these scores for the sample’s mutation rate, as our goal was to identify subtypes independent of mutation rate.

As reported previously by other groups [24], we observed large variations in the number of mutated cancer associated genes in each sample, ranging from −1,003, with a median of 91 mutated genes per sample (Figure 1A). We also observed differences between the cancer types, with a median of 2 mutated cancer-associated genes for chromophobe renal cell carcinoma (KICH) and papillary thyroid carcinoma (THCA), and 27 for pancreatic adenocarcinoma (PAAD). As expected, we observed a larger number of mutated pathways than of mutated cancer-associated genes across all samples (median = 103). Acute Myeloid Leukemia (LAML) had the lowest (12) and uterine corpus endometrial carcinoma (UCEC) the highest (238.5) median of mutated pathways.

**Figure 1.**
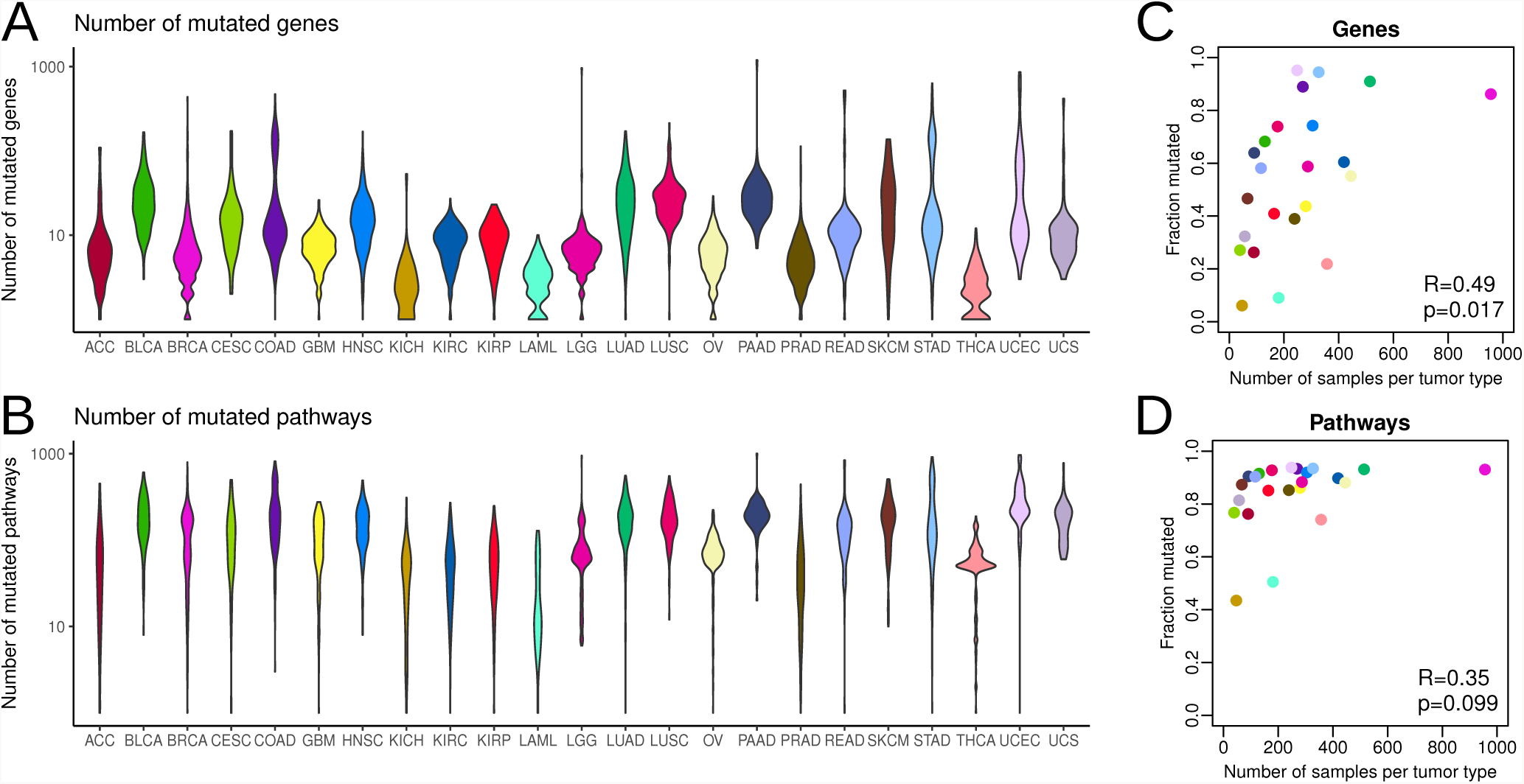
A–B) Violin plots visualizing the distribution of (A) the number of mutated genes and (B) the number of mutated pathways, for each of the 23 cancer types. C–D) The fraction of mutated genes (C) and pathways (D) plotted against the number of samples in each cancer type. R: Pearson’s correlation coefficient, p: Pearson’s correlation test p-value.

Apart from these differences, we did observe fairly similar numbers of pathways that were mutated in at least one sample in each of the 23 cancer types, indicating that the data were sufficiently de-sparsified. On average, 84.2% of all pathways were mutated in at least one sample of each cancer type, ranging from 43.4% in KICH to 93.8% in UCEC. These percentages were lower for mutations in cancer-associated genes, with an average of 54.8% of all cancer-associated genes being mutated in a cancer type (minimum of 6.1% for KICH, maximum of 95.1% for UCEC). While these numbers do depend on sample size for the gene mutation data (Pearson *R* = 0.49, *p* = 0.017, Figure 1C), the correlation of the fraction of mutated pathways with the number of samples available per cancer type is not significant (*p* = 0.099, Figure 1D). This again confirms that the data de-sparsification was successful (see also Supplemental Figure S2).

### Exhaustive search of dissimilarity metrics to inform subtype classification

We next wanted to identify a dissimilarity metric that would result in the best separation of the mutation data into subtypes. For each cancer type, we calculated the average distance between all patients on both the gene mutation and pathway mutation scores, using 12 dissimilarity metrics [16]. Binomial and Mahalanobis dissimilarities (see Supplemental Materials and Methods) best separated samples based on gene mutation scores, while the pathway mutation scores were best separated by the binomial dissimilarity index (Figure 2). We observed larger distances when using pathway mutation scores, with a median distance of 108 using binomial distance on pathway mutation scores compared to 21 when using Mahalanobis distance on gene mutation scores. This confirmed that de-sparsification of the data into pathway mutation scores helped in separating the samples. We therefore used the binomial distance on pathway mutation scores for the subtyping analyses we present in the following paragraphs.

**Figure 2.**
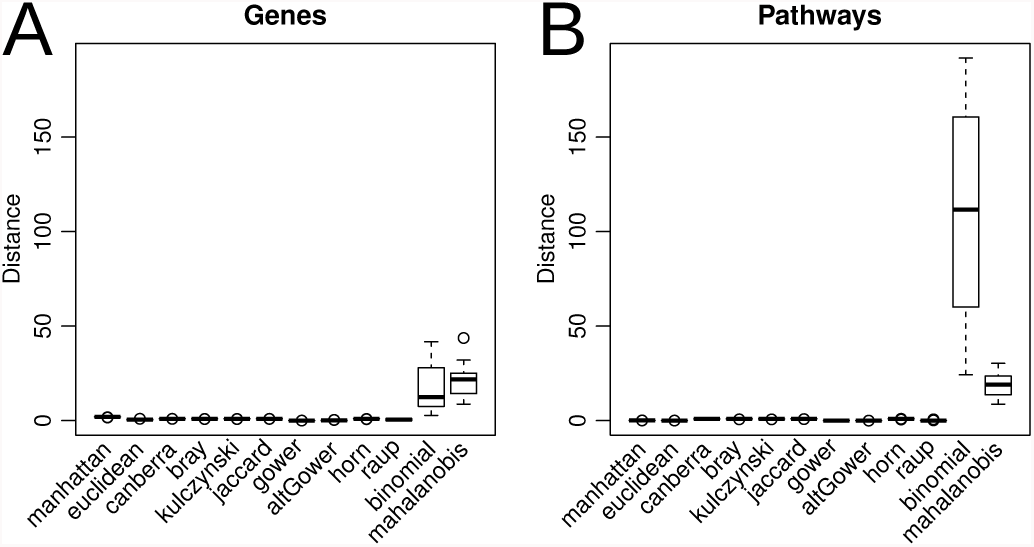
Boxplots of the median distance between samples using (A) gene mutation scores and (B) pathway mutation scores across the 23 cancer types, using different dissimilarity metrics. The line in the center of the box denotes the median, the box edges denote the first and third quartiles of the data, and the error bars denote 1.5*×* the interquartile range.

### Biological signatures encode histopathological information

We performed principal coordinate (PC) regression on the pathway mutation data to explore whether phenotypic or clinical information was retained in the most variable components of the data (see Methods). For each cancer, we reported the clinical parameters that were associated with the top five principal components (*p <* 0.01), and visualized these in a word cloud (Figure 3A). The most significant variables included age (nominally significant for 12/23 cancer types) followed by tumor grade (for 2/4 cancer types for which the variable was available) and histological type (6/13 cancer types). This was not unexpected—with age cells may build up somatic mutations, while high grade tumors divide faster, which may lead to more replication errors.

**Figure 3.**
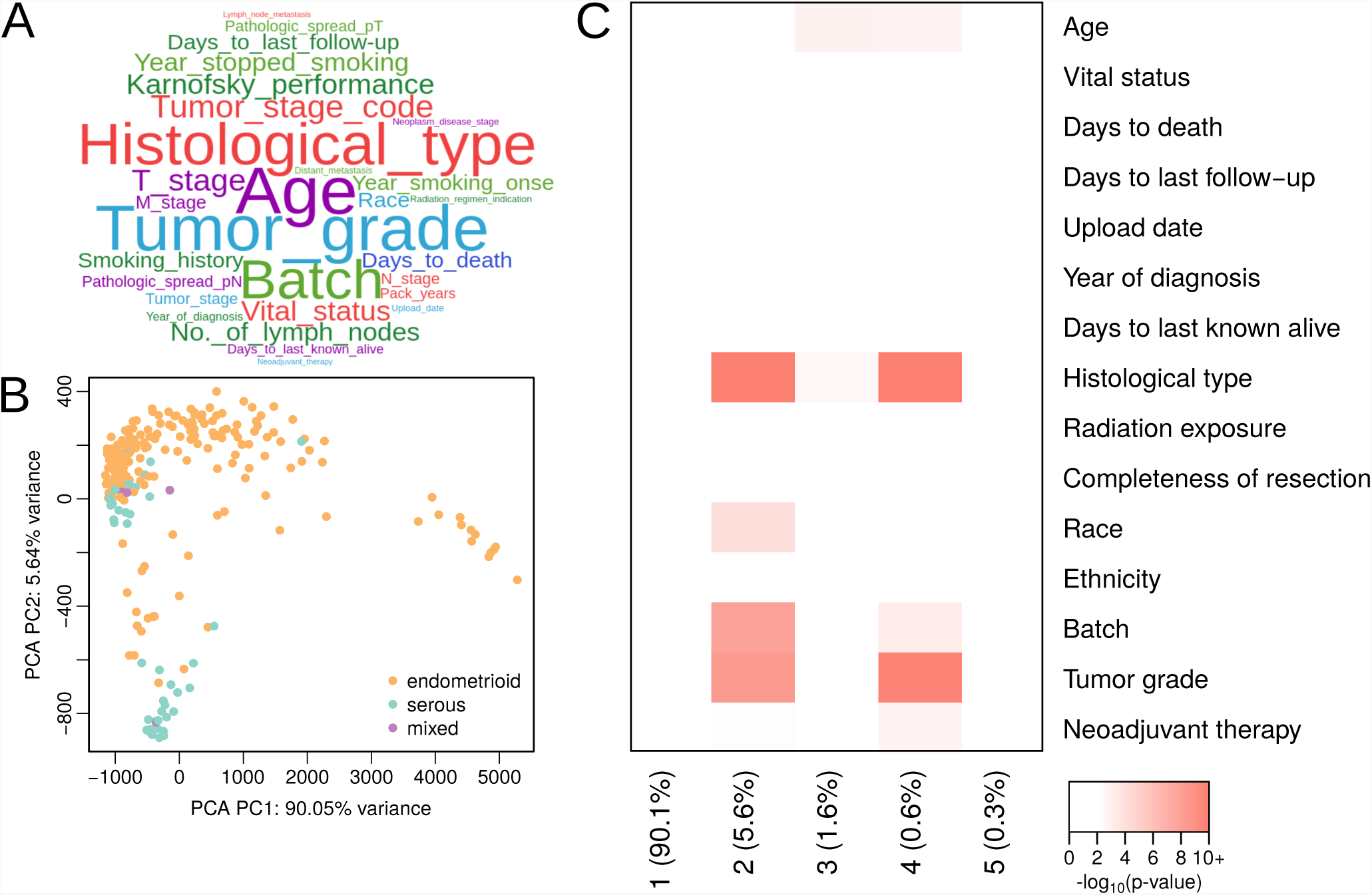
De-sparsified mutation data retains histological and other clinical information that might define cancer subtypes. A) Word cloud of the most associated variables with the top 5 principal components across the 23 cancer types. B) First two coordinates of the principal components for UCEC samples. Each point represents an individual sample. Colors represent the histological type of the cancer, orange being endometrioid endometrial adenocarcinoma, purple being mixed serous and endometrioid, and cyan representing serous endometrial adenocarcinoma samples. C) Heatmap of the −*log*_10_ p-values from PC regression against clinical variables.

In Figure 3B, we show the Principal Component Analysis (PCA) plot for UCEC, colored by histological type. We observe a separation of serous endometrial adenocarcinoma samples from most endometrioid endometrial adenocarcinoma samples. Heatmaps of the PC regression helps inform this particular visualization, as we see several associations for the top 5 components and various phenotypic and clinical variables in the UCEC samples (Figure 3C). The first principal component, which explains most of the variance in the data, did not significantly associate with any clinical variable in UCEC. However, after inspecting the heatmaps in Supplemental Figure S2C–D, we thought this component might associate with a sample’s mutation rate. To test this, we investigated the mutation rates of the 13 samples that cluster separately from the rest, and found significantly higher mutation rates in these tumors (t-statistic = 13.4, p-value = 1.27. 10^*−8*^). This means that, while we correct for a sample’s mutation rate, it is not completely filtered out, and still dominates some of the clustering in UCEC. However, in our subtyping analyses described below, we further corrected for mutation rate by removing samples that clustered separately (*<* 10 samples in the tumor-specific subtyping analysis, and *<* 50 samples in the pan-cancer analysis) from most other samples.

After correcting for multiple testing, we identified a number of statistically significant (adjusted *p <* 0.05) variables (Supplemental Table 2). Multiple variables, including age, histological type, race, and variables associated with follow-up and patient survival were significant in several tumor types. Additionally, technical variability (batch number) appears to associate with pathway mutation scores in a number of cancers. We also identified variables that were relevant for specific cancer types, including smoking history and “years stopped smoking” for lung adenocarcinoma (LUAD), and Gleason score for prostate adenocarcinoma (PRAD).

### Identification of prognostic mutation subtypes

We explored whether we could identify subtypes associated with cancer survival. We used the binomial dissimilarity index to cluster the de-sparsified mutation data of each cancer type. We split the cluster dendrograms in 2 − 4 groups (see Methods). We identified significant prognostic subtypes (log-rank test *p <* 0.01) for three cancer types—adrenocortical carcinoma (ACC), LAML, and low-grade glioma (LGG). Pathways associated with these prognostic subtypes are listed in Supplemental Table 3.

Clustering ACC samples in two groups produced subtypes of 74 and 12 patients. The smaller subtype was associated with poor survival (log-rank test *p* = 0.0027, Figure 4A). Twenty-two pathways were associated with this subtype (see Methods), including pathways involved in apoptosis and cell cycle, both of which are known hallmarks of cancer [25], and Wnt and Notch signaling, both known cancer drivers. Protein expression levels of Notch pathway genes have previously been associated with clinical outcome in ACC [26] and Wnt signaling has been reported to play a role in differentiation of the zone glomerulosa of the adrenal cortex [27]. While Wnt/*β*-catenin signaling has been reported by the TCGA as frequently altered in ACC [28], this pathway had not previously been reported to be associated with survival. In addition to these known cancer driver pathways, we identified mutations in neurotrophin signaling, which plays a role in neuron development and differentiation [29], in the poor prognosis subtype.

**Figure 4.**
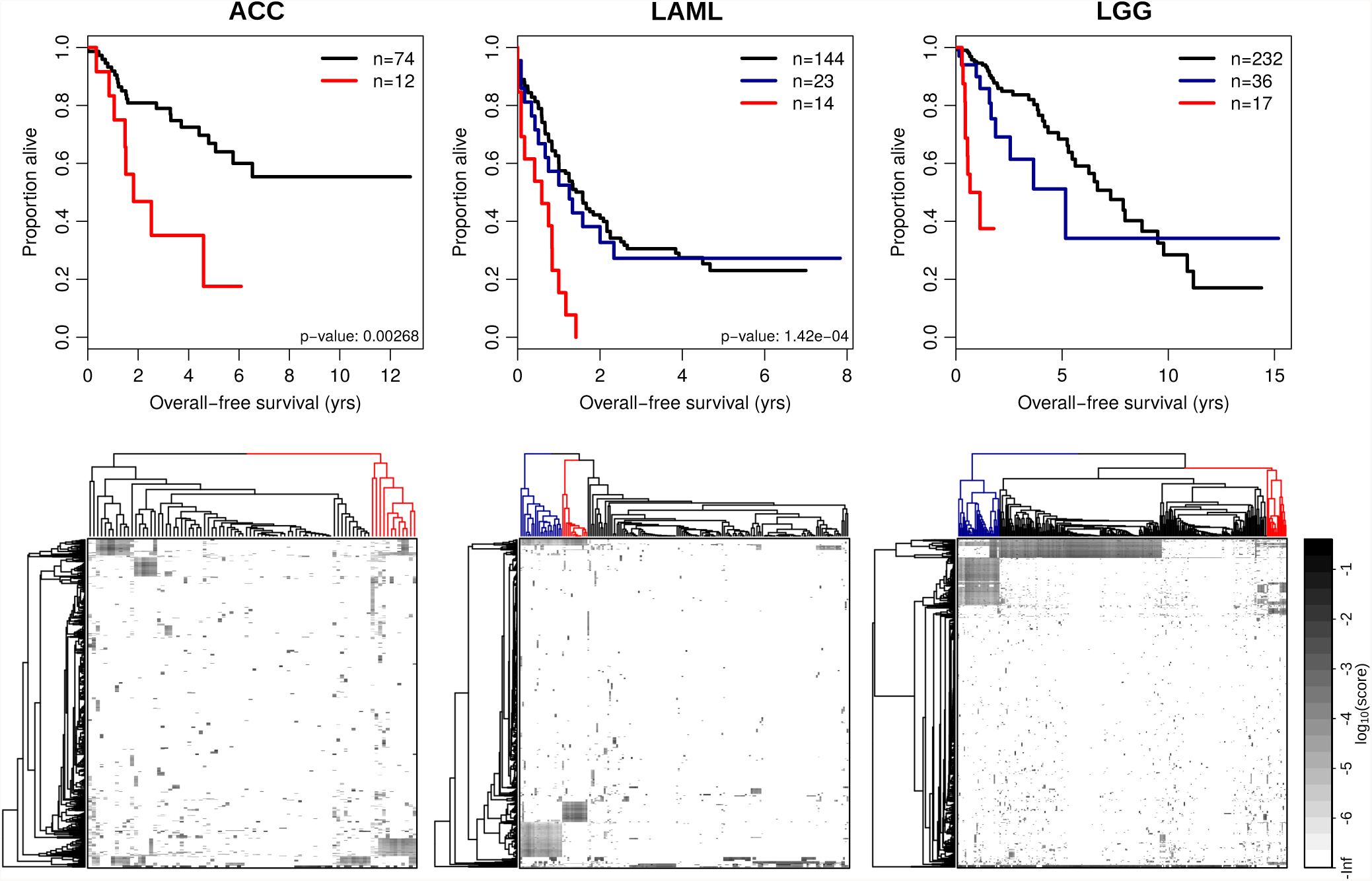
A) Kaplan-Meier plots depicting overall survival for patients with different prognostic subtypes in ACC, LAML, and LGG. Overall survival of patients in the poor prognostic group is shown in red. Plots include the number of samples in each subtype, as well as the log-rank test p-values. B) Heatmaps of pathway mutation scores (shown on a log-scale). The subtypes are visualized in the column dendrograms, with the same color coding as used in (A).

Splitting the LAML samples in three clusters produced a poor survival subtype of 14 patients (log-rank test *p* = 1.4⋅10^−^4), compared to two larger clusters including 23 and 144 patients (Figure 4B). Thirty-eight pathways were associated with this poor survival cluster, of which 11, with roles in apoptosis, cell cycle, and Notch and Wnt signaling, were a subset of the pathways we identified in the poor survival subtype of ACC. Expression of both Notch and Wnt signaling genes has been implicated in LAML [30, 31], but mutational patterns of these pathways had not been reported to be associated with patient survival. In addition to these pathways, DNA damage response pathways, which included p53 and ATM signaling, were mutated in the poor prognostic subtype of LAML.

Finally, splitting the LGG data in three clusters resulted in a subtype of 232 patients, a subtype with somewhat worse prognosis including 36 patients, and a poor prognosis subtype of 17 patients (log-rank test *p* = 1.3 · 10^−13^, Figure 4C). Tumors assigned to this subtype had higher mutation scores in multiple EGF receptor family pathways, in cell-cell contact and cellular structure (“adherens junction,” “gap junction”), the immune system (“cytokine-cytokine receptor interaction”), and in brain tissue-associated pathways (including “gonadotropin-releasing hormone signaling”). Mutations in cell-cell contact genes could be important for metastasis, and immune cells are known to play a critical role into transforming low grade glioma into glioblastoma [32]. *EGFR* is a known glioma driver [33]. However, while this gene was mutated in 15/17 patients, other genes belonging to EGF receptor family pathways, including *EGF*, *GNAS*, and *PTRB*, were also mutated in tumors belonging to this subtype, and might not have been detected if we had focused on *EGFR* alone.

Because the sample sizes of the poor prognostic subtypes were small in comparison to the subtypes with relatively better prognosis, we wanted to make sure these subtypes were not identified due to random selection of a small group of patients with poor prognosis. For each of the significant cancer types we performed 10, 000 sample label permutations and found that the original log-rank tests were more significant than those on the permuted null background (Benjamini-Hochberg adjusted p-values *<* 0.01), indicating that the poor prognostic subtypes were not identified by chance, but were detected based on mutations in specific biological pathways (see Supplemental Figure S3). We would like to note that, even though we did not identify any associations in the permuted data, the poor prognosis subtypes could still be confounded with specific clinical characteristics. We assessed whether the LGG subtype was confounded by indication of radiation therapy, but did not find a significant association (Chi-squared test p-value = 0.123). For ACC and LAML, unfortunately, no detailed information on treatment status was available. While the numbers of patients in the identified poor prognosis subtypes are small, and while these subtypes might be associated with certain patient characteristics, we believe that treating these patients with personalized therapy regimens that specifically target the subtype-specific signaling pathways could dramatically improve prognosis in these patients.

### Integration of cancer type-specific subtypes with drug targeting information

While somatic mutations have been analyzed in the context of actionable drug targets [34], to our knowledge, mutation subtypes of multiple cancer types have not been integrated with drug targeting databases. To identify therapies targeting pathways that are specifically mutated in cancer subtypes, we performed an enrichment analysis using the Connectivity Map (CMap) [18]. In short, we identified subtypes using the analysis described above, identified pathways that were frequently mutated in these subtypes, and selected genes belonging to these pathways that were more frequently mutated in the subtype of interest. We matched these genes against CMap to identify drugs with targets enriched for these genes (see Methods). We identified 251 subtype-drug interactions for a total of 15/23 cancer types. For each cancer type, we selected interactions with the highest enrichment scores and reported these in Table 1.

For the poor prognostic subtype in ACC that we described above, we observed an enrichment of targets of MDM2 inhibitors (idasanutlin and nutlin-3). This was not unexpected, as we found specific mutations in p53 pathways in this subtype. We did not identify any enriched drug targets for the poor prognosis subtypes of LAML and LGG. However, only two Notch inhibitors are available in CMap, and both are listed without any targets. While many EGFR inhibitors are available in CMap, *EGFR* was the only gene from the EGFR pathways that overlapped with EGFR drug targets. Since we only considered drug-subtype interactions for subtypes with at least two mutated drug targets, these drugs were not included in our analysis. However, the subtypes with intermediate survival were enriched for lonafarnib—a farnesyltransferase inhibitor—in LAML, and atiprimod— an angiogenesis inhibitor—and JAK inhibitors in LGG. Lonafarnib was also enriched in a large subtype of THCA tumors, while JAK inhibitors were enriched in BLCA, HNSC, KIRC, and LUSC. In addition to these drugs, we identified multiple receptor tyrosine kinase inhibitors associated with subtypes in BLCA, GMB, and HNSC. We also identified a PI3K inhibitor (wortmannin) to be enriched in mutated targets in a UCS subtype, a proteasome inhibitor (bortezomib) in KIRP and OV subtypes, and a CREBBP/EP300 inhibitor (SGC-CBP30), in a small subgroup of BRCA and PRAD patients.

In total, the subtypes that were enriched for mutations in drug targets from CMap account for 12% (689/5805) of all primary tumors from TCGA. This is a substantial number of patients, considering the strict thresholds we used for this analysis. We believe that, as more drug target information becomes available, we may find additional subtype-drug associations that might help identify subgroups of patients that could benefit from targeted treatment of their tumor’s mutational profile. This also suggests that a pathway-based analysis might provide a window into new therapeutic options for cancer patients.

### A pan-cancer analysis identifies 9 mutation subtypes

Finally, using the clustering technique described above, we performed a subtyping analysis across all 23 cancer types. Because of the large number of samples (*n* = 5, 805), and because the number of potential pan-cancer mutation subtypes is unknown, we divided the cluster dendrogram into *k* subtypes, ranging *k* from 2 to 1, 000, removing clusters of size *<* 50. We selected the largest *k* for which we observed a “breakpoint” in the sample size of the largest cluster (Supplemental Figure S4) that had ≥ 90% of all samples assigned to a subtype (belonged to a cluster of size ≥ 50). This returned 9 pan-cancer subtypes, with an average sample size of 581, ranging from 74 (subtype S8) to 2, 194 (S5) (Figure 5A–B and Supplemental Figure S5A).

**Figure 5.**
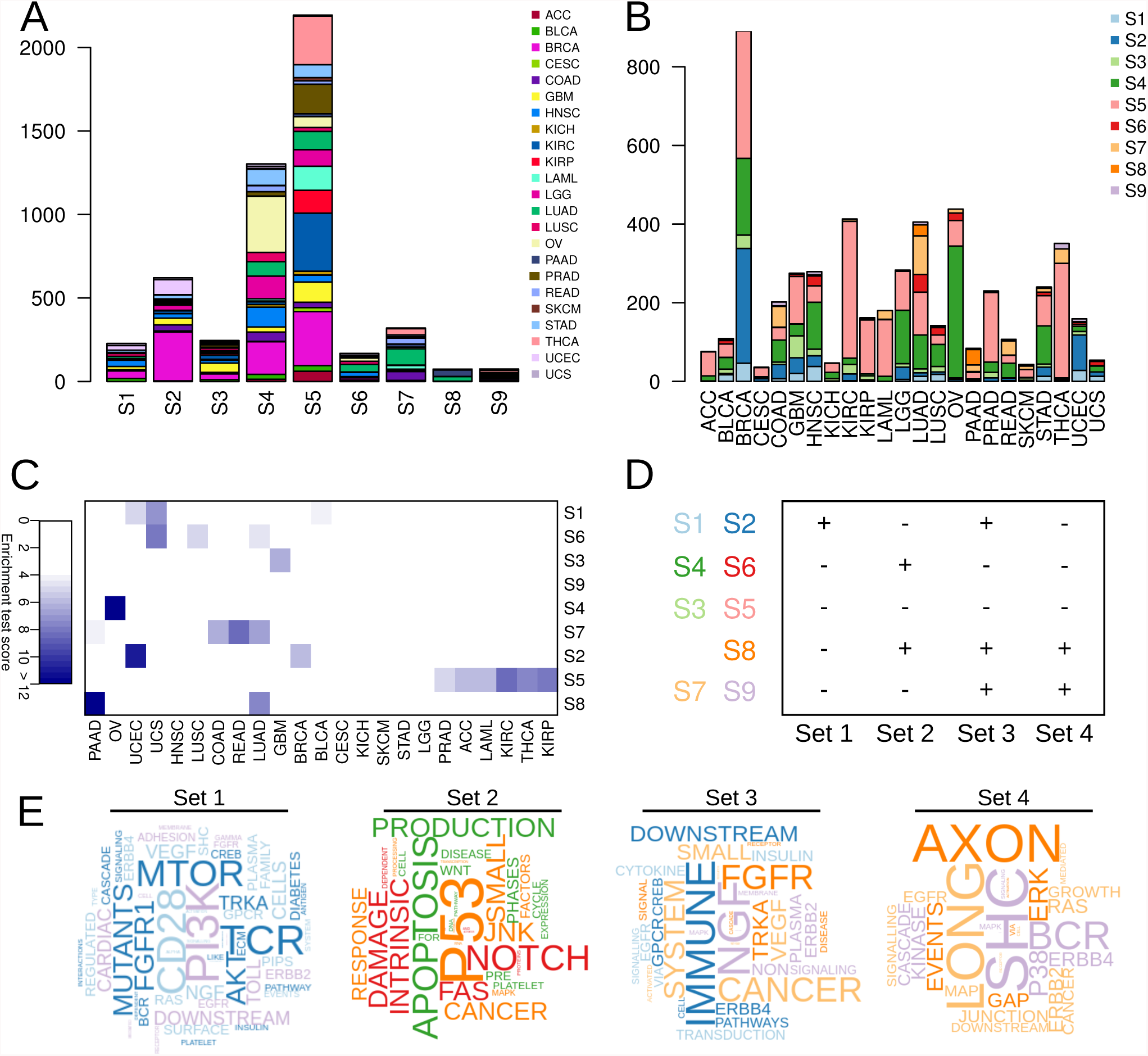
Pan-cancer mutation subtyping results. Stacked bar charts of A) the number of the different cancer types in each of the 9 subtypes, B) the number of the different subtypes per cancer type. C) Enrichment of subtypes for specific cancer types. Test scores are calculated by multiplying the odds ratio to 1 for significant results (Bonferroni-adjusted *p <* 0.05), and to 0 for nonsignificant results. Significant scores with odds ratio’s *>* 4 are visualized in blue. D) Four overarching sets of mutations are associated (+) with multiple subtypes. E) Word clouds visualize enrichment for specific words associated with each of the four sets of pan-cancer pathways.

The nine pan-cancer subtypes were all heterogeneous with respect to cancer type—each subtype included samples from at least 7/23 and on average 18/23 different cancer types (Figure 5A). In addition, all cancer types were represented in at least 4/9 and on average in 7/9 subtypes (Figure 5B). Some cancer types were over-represented in particular subtypes (Figure 5C). For example, S1 was enriched for UCEC, uterine carcinosarcoma (UCS), and BLCA, which are anatomically close cancers. S2 was enriched for UCEC and BRCA, both cancers associated with the female reproductive system. S5, the largest of the nine subtypes, was enriched for ACC, kidney renal clear cell carcinoma (KIRC), kidney renal papillary cell carcinoma (KIRP), LAML, PRAD, and THCA. S6 was enriched for lung cancers (LUAD, lung squamous cell carcinoma (LUSC)), as well as UCS. S7 was enriched for four types of adenocarcinoma—colon adenocarcinoma (COAD), LUAD, PAAD, and rectum adenocarcinoma (READ), and S8 was enriched for LUAD and PAAD.

To identify whether these nine pan-cancer mutation subtypes associated with a high mutation rate in specific biological pathways, we selected pathways with mutations in *>* 95% of all samples in a specific subtype. This resulted in a total of 202 pathways (Supplemental Figure S5B). All of the subtypes, except S5, which had no subtype-specific pathway mutations, had frequent mutations in Kyoto Encyclopedia of Genes and Genomes (KEGG) “pathways in cancer.” Some subtypes (S1 and S2, S4 and S6) had exactly the same sets of pathways mutated, but with different average mutation rates. Higher mutation rates were observed in S2 and S4 compared to S1 and S6, respectively (t-statistic= 5.66, *p* = 3.56 ⋅ 10^−8^ for S2 versus S1, t-statistic= 3.10, *p* = 2.73 ⋅ 10^−3^ for S4 versus S6). This may indicate that patients belonging to these subtypes might benefit from (additional) immunotherapy.

Using hierarchical clustering (binomial distance) on average mutation scores in these 202 pathways, we identified four overarching “sets” of pathways that were differentially mutated in the pan-cancer subtypes (see Figure 5D–E and Supplemental Figure S5B). The first set of pathways was highly frequent in subtypes S1 and S2, and was enriched for PI3K/Akt/mTOR signaling and immune system pathways (including T-cell receptor and CD28 signaling). The second set was frequent in S4, S6, and S8, and involved DNA damage pathways, apoptosis, and Notch signaling. The third set was highly mutated in 5/9 subtypes (S1–2, S7–9), and included immune system and metabolism pathways and several growth factor receptor pathways (FGFR, ERBB2/4, VEGF). Finally, the fourth set of pathways was highly mutated in S7– 9, and included neuronal pathways, signaling via SHC family adapter proteins, B-cell receptor pathways, and pathways related to cell-cell contacts.

These four sets of pathways highlight important processes that are highly recurrently mutated in large sub-groups of patients, across different cancer types. Interestingly, there does not seem to be a dependency between these four sets of highly recurrent processes. Most of the sets are either mutated alone (such as Set 2 in pan-cancer subtypes S4 and S6) or in combination with other sets (for example Sets 2–4 in pan-cancer subtype S8). An exception to this is Set 1, which does not co-occur with Sets 2 and 4, indicating a possible mutual exclusive relationship between PI3K/Akt/mTOR on the one hand, and p53 and certain growth factor receptor pathways on the other. These results could indicate that patients assigned to certain subtypes may benefit from targeting specific pathways in selecting therapies, while others may need combinations of targeted treatment approaches to target multiple processes that are disrupted. Finally, subtypes S3 and S5 do not have any recurrently (*>* 95%) mutated pathways, indicating that treating these patients may require individualized analysis of their unique mutational patterns in deciding on a precision medicine strategy.

### Pan-cancer mutational patterns correspond to pathway activation levels and response to drug inhibition

We validated the pan-cancer subtypes we identified in two ways. First, we wanted to make sure that the signaling pathways we had identified as highly mutated in the different pan-cancer subtypes were active. To do this, we integrated these results with Reverse Phase Protein Array (RPPA) data from TCGA. We calculated pathway activation scores for Akt signaling and DNA damage response pathways, corresponding to the “Set 1” and “Set 2” pathways we had identified in Figure 5E (see Methods). We did not calculate activation scores for Sets 3– 4, because the RPPA data did not include enough proteins corresponding to genes that belonged to these sets of pathways. We identified significantly higher protein activation scores in tumors from patients belonging to the subtypes that had higher levels of mutations in these pathways. We identified a mean Akt pathway protein activation score of 0.459 in patients belonging to “Set 1” subtypes (S1–2), compared to a mean score of 0.0756 in other patients (two-sample t-test t-statistic = 2.34, *p* = 0.0198), and a mean DNA damage response protein activation score of 0.283 in patients belonging to “Set 2” subtypes (S4, S6, S8) compared to 0.158 in other patients (t-statistic = 2.45, *p* = 0.0146). This indicates that the pan-cancer subtypes we had identified based on pathway mutation scores also corresponded to higher protein levels in these pathways.

Second, we wanted to determine whether cell lines with mutations in the overarching sets of pathways we had identified in the pan-cancer subtypes were more sensitive to drugs targeting those pathways. We downloaded mutation and drug response data from the Cancer Genome Project (CGP). Again, we focused on pathways identified in Sets 1–2, for which drug targeting information was available in CGP. We identified which cell lines had mutations in all “Set 1” pathways, and compared how these cells responded to PI3K/MTOR inhibitors compared to other cell lines (see Methods). We identified significant differences (FDR *<* 0.05) in response to 5/24 PI3K/MTOR inhibitors. Most of which (4/5) had significantly lower IC50s (median t-statistic = −3.13, largest effect observed for Pictilisib), indicating that these cell lines were more responsive to PI3K/MTOR inhibition (see also Supplemental Table 4). We repeated this analysis for cell lines with mutations in all “Set 2” pathways and compared how these cell lines responded to drugs interfering with DNA replication. We identified significant differences for 7/11 of these drugs, all of which showed significantly higher IC50s (median t-statistic = 4.401), indicating that these cell lines were less responsive to drugs interfering with DNA replication (see also Supplemental Table 4). While this result may seem counterintuitive at first, it is known that tumors with impaired DNA damage response may become resistant to chemotherapy [35].

In summary, by identifying pan-cancer subtypes, we were able to uncover processes that play a role in large subgroups of cancer patients, that are (to a degree) independent of cancer type. These subgroups indicate that core pathways are often mutated, independent of the tissue of origin and may indicate specific targets for therapeutic intervention that should be explored.

## DISCUSSION

Even though exome sequencing data is now available for large numbers of tumors, identifying mutation subtypes in cancer is still a challenge due to the sparseness and heterogeneity of the data. We developed SAMBAR, a de-sparsification method that summarizes somatic mutations in genes into pathway level mutation scores. We showed that SAMBAR helps identifying mutational patterns associated with clinical phenotypes and prognosis, potential targeted treatment options for cancer-specific subtypes, as well as mutational patterns that are manifested across multiple cancer types.

Some of the pathways we identified in the prognostic subtypes, including cell cycle, apoptosis, and DNA damage response, are frequently mutated in cancer. However, subgroups of patients may still benefit from specific targeted treatment options that can be found through a pathway-level analysis and through developing methods to interrogate diverse data resources. For example, we identified MDM2 inhibitors as potential targets for treatment of the poor prognosis subtype in ACC by integrating subtype-specific mutations with a drug targeting database. In addition, several signal transduction pathways for which targeted treatment options are available are specifically mutated in the poor prognosis subtypes we identified. These pathways include Notch signaling in ACC and LAML and EGF receptor family pathways in LGG. Thus, by considering mutations in all of the genes associated with subtype-specific mutation of signaling pathways, we may find additional patients that could benefit from targeted treatment options.

The results from this analysis suggest that, rather than only focusing on well known “driver” genes, we should also include genes associated with particular biological pathways when profiling patients to search for personalized treatment options. By generating patient-specific “pathway mutation profiles,” we may not only identify patients who could benefit from specific targeted therapeutics, but we will also obtain a clearer picture of the cellular processes that are altered through mutation in a specific tumor. This expanded pathway- and process-based approach may help identify combination therapies that target multiple pathways that are altered in a patient’s tumor.

In our pan-cancer analysis, we identified four overarching types of mutational patterns. The first set included PI3K/Akt/mTOR signaling pathways. This means that, likely, a large number of patients will benefit from targeted inhibition of this signal transduction pathway. In addition, one of the pan-cancer mutational patterns we identified was enriched for several growth factor pathways, including EGF receptor family genes, and FGFR and NGF signaling. Targeted treatment options are available for each of these pathways. Because these pathways are mutated in a large set of the primary tumors we analyzed, we believe that these treatment options are worthy of further investigation and may lead to better treatment options for a large numbers of patients.

Recently, large hospitals have started to include mutational profiling as a standard procedure to characterize tumors, and to assign patients to available targeted treatment options or ongoing clinical trials targeting specific mutations [36]. However, therapeutic decisions are typically tied to examining single gene mutations. Our framework to classify cancers based on mutational patterns in biological pathways could help expand precision medicine applications by both identifying groups of patients who may or may not respond to particular therapies and by identifying pathways that might be useful targets for therapeutic intervention.

Finally, while we concentrated on identifying subtypes that include relatively large numbers of patients, the pace at which we are collecting mutational data continues to accelerate. As more data become available, we can fine-tune our subtyping analysis, identifying not only the largest, but also smaller groups of patients for which targeted treatment options may be available. Analysis of larger sample sizes will also improve as our understanding of the biological pathways that are important in driving cancer. Future research may focus on combining our pathway mutation scores with previously published network propagation methods to further fine-tune classification of pathway mutation profiles.

## ACKNOWLEDGMENTS

We would like to thank Kimberly Glass, PhD and Alessandro Marin, PhD, as well as all members of the Quackenbush laboratory for helpful suggestions on the manuscript.

## AUTHOR CONTRIBUTIONS

Conceptualization, M.L.K, W.D., J.Q.; Methodology, M.L.K, J.N.P.; Formal Analysis, M.L.K., J.N.P.; Investigation, M.L.K., J.N.P; Resources, J.Q., W.D.; Data Curation, M.L.K., P.S.; Writing–Original Draft, M.L.K., J.N.P; Writing–Review & Editing, M.L.K., J.N.P., P.S., W.D., J.Q.; Visualization, M.L.K., J.N.P.; Supervision, W.D., J.Q.; Funding Acquisition, M.L.K., W.D., J.Q.

## SUPPLEMENTAL MATERIALS AND METHODS

### Curation of clinical data

We used R package RTCGAToolbox to download clinical data for all TCGA. We used the *getFirehoseDatasets()* function to get a list of all tumors (23 when we accessed these data), and the function *getFirehoseRunningDates()* to get a list of the available Firehose dates. We prepared the clinical data for each cancer type separately. First, we checked at which of the available Firehose dates clinical data had been uploaded. We then downloaded clinical data from the first available Firehose date that included this data type for the cancer type of interest, as well as for the next available date. We made column names consistent by removing all punctuation characters and spaces from these names, and by converting all characters to lowercase. We then merged the data by keeping all data from the latest (of the two) Firehose dates, but added any patients or columns from the previous date that were not available in the data uploaded at the later date. We continued this, comparing clinical data uploaded at a particular time point to the combined set of data previously uploaded. This way, we contained as much clinical information as possible, while containing the most up-to-date entries for those data points that were uploaded multiple times (for example, follow-up data). For visualization of clinical parameters in Figure 3, we converted phenotypic parameters according to Supplemental Table 1. We merged duplicate clinical parameters, such as “years to birth” and “age at diagnosis.”

### Dissimilarity metrics

The Mahalanobis distance is similar to Euclidean distance except that it normalizes the data based on a covariance matrix, making the distance metric scale-invariant. Intuitively, it asks what the distance of a point in n-dimensional space is for a given n-space distribution. The binomial dissimilarity index is derived from the binomial deviance under the null hypothesis that the two compared communities are equal. For our data, this implies that the proportions of various signatures are more similar amongst similar samples.

The larger the distances or dissimilarity measures between the samples of a specific cancer, the greater the hidden structure revealed to help in differentiating samples potentially linked to phenotypic properties. The greatest median measurement and variation amongst cancer types when using the binomial dissimilarity index on the biological pathway de-sparsified mutation data, and therefore selected this metric to identify cancer-specific mutation subtypes associated with survival.

---

**Supplemental Figure S1.**
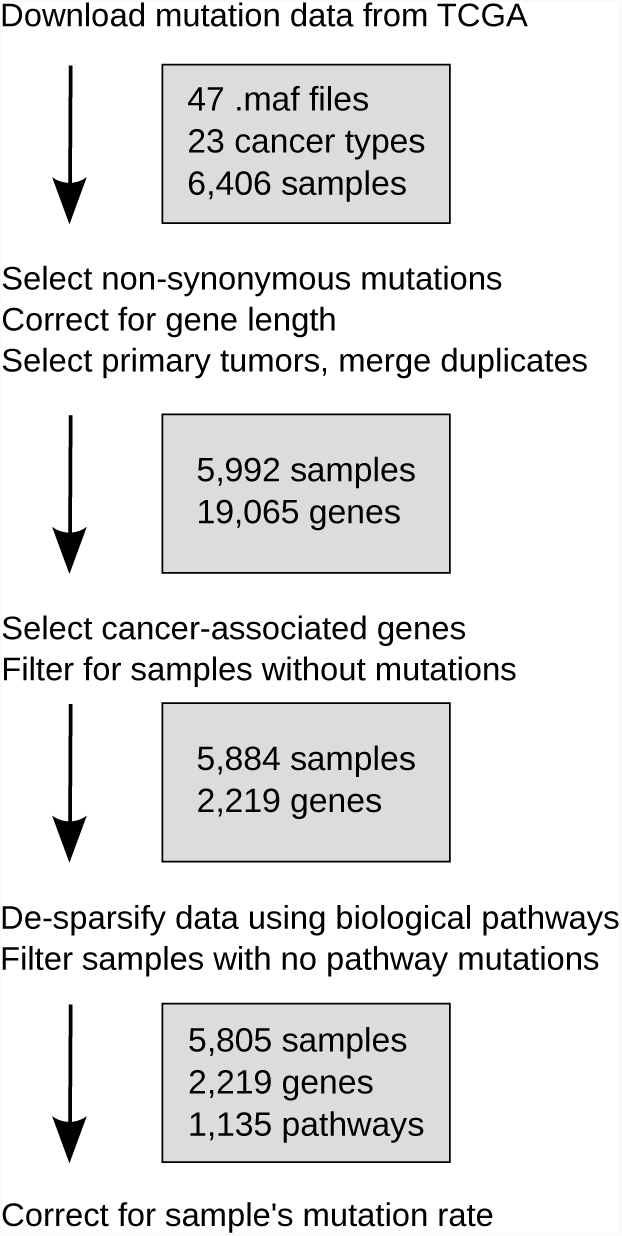
Mutation data pre-processing workflow.

**Supplemental Figure S2.**
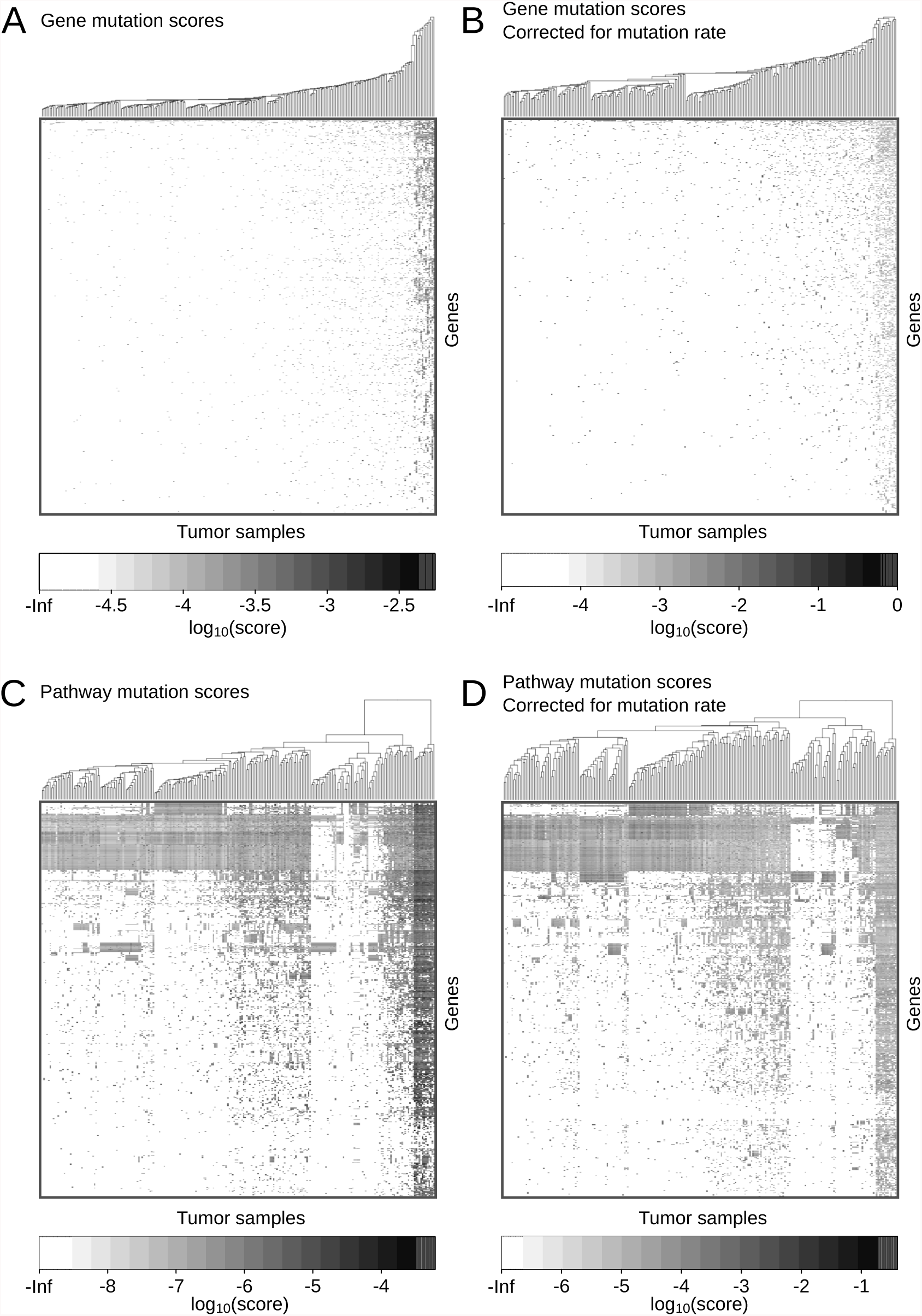
Hierarchical clustering (Euclidian distance) of 246 uterine corpus endometrial carcinoma (UCEC) samples based on A) gene mutation scores, B) mutation scores corrected for each sample’s mutation rate, C) pathway mutation scores, and D) pathway mutation scores corrected for each sample’s mutation rate. Heatmaps visualize the −*log*_10_ of the mutation scores. For visualization purposes, mutation scores of 0 were set to the lowest −*log*_10_ non-zero mutation rate across all samples in each specific data type.

**Supplemental Figure S3.**
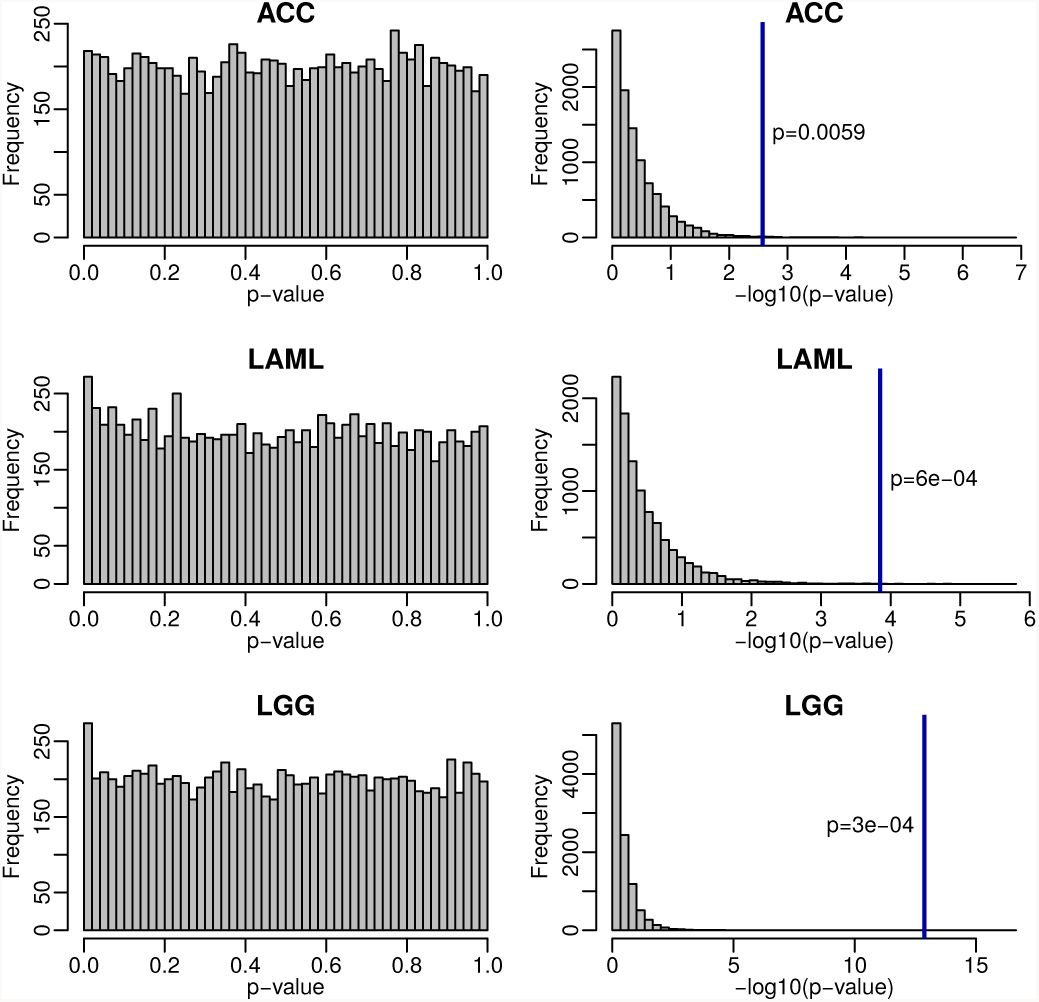
Histograms of p-values (left plots) and -*log*_10_ p-values (right plots) of 10, 000 log-rank tests with permutated sample labels. Permutations are based on group sizes and clinical data of the cancer types for which we identified significant prognostic subtypes (log-rank test *p <* 0.05, ACC, LAML, and LGG). The blue lines in the plots on the right illustrate the -*log*_10_ p-values which we observed from the log-rank test on the non-permuted data (as in Figure 4). The p-value next to the blue lines is the Bonferroni-adjusted permutation p-value.

**Supplemental Figure S4.**
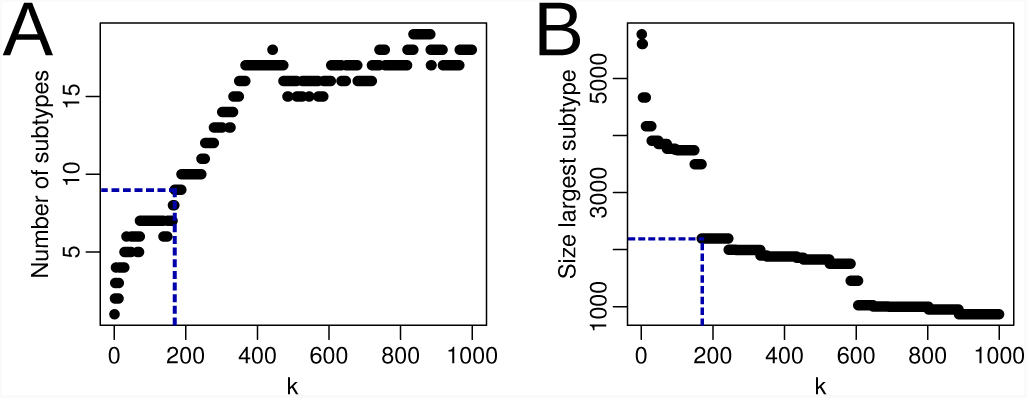
A) Number of subtypes identified and B) size of the largest subtype at different thresholds of *k*. The blue dashed line indicates *k* = 169.

**Supplemental Figure S5.**
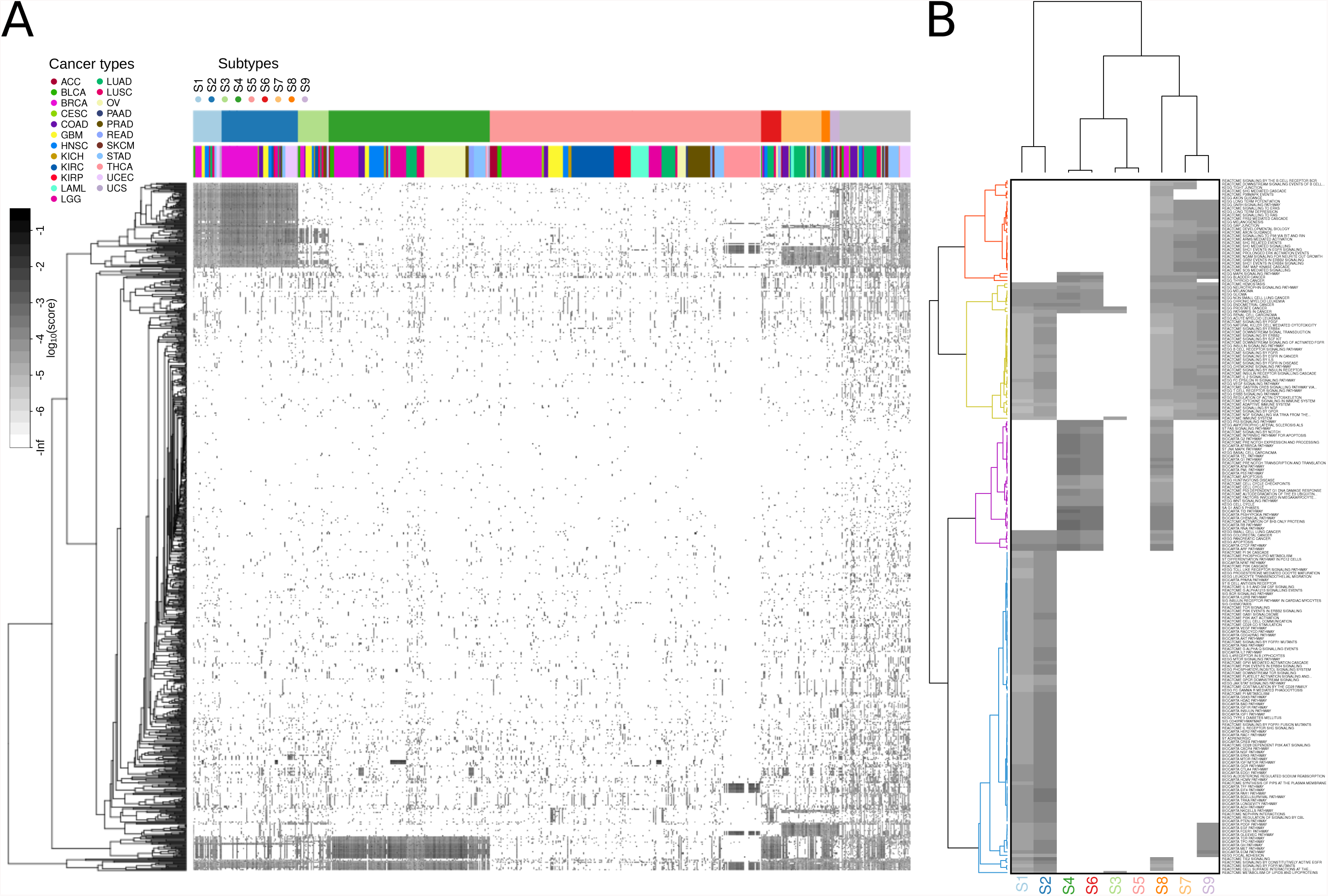
A) Heatmap depicting the de-sparsified mutation data. Rows (pathways) are clustered using binomial distance on pathway mutation scores. Columns are ordered based on the subtype a sample belongs to, and the tumor type a sample was harvested from. The column color bars depict to the pan-cancer subtypes (top) and cancer types (bottom). B) Heatmap depicting pathways with mutations in *>* 95% of all samples of one or multiple subtypes. Rows and columns are clustered using binomial distance. The set of 202 frequently mutated pathways cluster into four main groups, which are visualized with different colors in the row dendrogram. Pathway mutation scores are shown on the same scale as in Supplemental Figure S2D.

## SUPPLEMENTAL TABLE LEGENDS

Supplemental tables are available online.

**Supplemental Table 1.** Table listing the clinical features from Firehose (processed as described in Supplemental Materials and Methods) and the feature names we assigned to them for visualization purposes.

**Supplemental Table 2.** Phenotypic variables that statistically associate (adjusted *p <* 0.05) with one of the first five principal components of the pathway mutation scores of a given cancer type.

**Supplemental Table 3.** Table listing pathways with poor prognosis signatures in ACC, LAML, and LGG.

**Supplemental Table 4.** Table listing results of the drug response analyses on cell lines from the Cancer Genome Project (CGP).

